# The landscape of MHC-presented phosphopeptides yields actionable shared tumor antigens for cancer immunotherapy across multiple HLA alleles

**DOI:** 10.1101/2023.02.08.527552

**Authors:** Zaki Molvi, Martin G. Klatt, Tao Dao, Jessica Urraca, David A. Scheinberg, Richard J. O’Reilly

## Abstract

**Background:** Certain phosphorylated peptides are differentially presented by MHC molecules on cancer cells characterized by aberrant phosphorylation. Phosphopeptides presented in complex with the human leukocyte antigen HLA-A*02:01 provide a stability advantage over their nonphosphorylated counterparts. This stability is thought to contribute to enhanced immunogenicity. Whether tumor-associated phosphopeptides presented by other common alleles exhibit immunogenicity and structural characteristics similar to those presented by A*02:01 is unclear. Therefore, we determined the identity, structural features, and immunogenicity of phosphopeptides presented by the prevalent alleles HLA-A*03:01, -A*11:01, -C*07:01, and - C*07:02.

**Methods:** We isolated peptide-MHC complexes by immunoprecipitation from 10 healthy and neoplastic tissue samples using mass spectrometry, and then combined the resulting data with public immunopeptidomics datasets to assemble a curated set of phosphopeptides presented by 20 distinct healthy and neoplastic tissue types. We determined the biochemical features of selected phosphopeptides by in vitro binding assays and in silico docking, and their immunogenicity by analyzing healthy donor T cells for phosphopeptide-specific multimer binding and cytokine production.

**Results:** We identified a subset of phosphopeptides presented by HLA-A*03:01, A*11:01, C*07:01 and C*07:02 on multiple tumor types, particularly lymphomas and leukemias, but not healthy tissues. These phosphopeptides are products of genes essential to lymphoma and leukemia survival. The presented phosphopeptides generally exhibited similar or worse binding to A*03:01 than their nonphosphorylated counterparts. HLA-C*07:01 generally presented phosphopeptides but not their unmodified counterparts. Phosphopeptide binding to HLA-C*07:01 was dependent on B-pocket interactions that were absent in HLA-C*07:02. While HLA-A*02:01 and -A*11:01 phosphopeptide-specific T cells could be readily detected in an autologous setting even when the nonphosphorylated peptide was co-presented, HLA-A*03:01 or -C*07:01 phosphopeptides were repeatedly nonimmunogenic, requiring use of allogeneic T cells to induce phosphopeptide-specific T cells.

**Conclusions:** Phosphopeptides presented by multiple alleles that are differentially expressed on tumors constitute tumor-specific antigens that could be targeted for cancer immunotherapy, but the immunogenicity of such phosphopeptides is not a general feature. In particular, phosphopeptides presented by HLA-A*02:01 and A*11:01 exhibit consistent immunogenicity, while phosphopeptides presented by HLA-A*03:01 and C*07:01, although appropriately presented, are not immunogenic. Thus, to address an expanded patient population, phosphopeptide-targeted immunotherapies should be wary of allele-specific differences.

**What is already known on this topic –** Phosphorylated peptides presented by the common HLA alleles A*02:01 and B*07:02 are differentially expressed by multiple tumor types, exhibit structural fitness due to phosphorylation, and are targets of healthy donor T cell surveillance, but it is not clear, however, whether such features apply to phosphopeptides presented by other common HLA alleles.

**What this study adds –** We investigated the tumor presentation, binding, structural features, and immunogenicity of phosphopeptides to the prevalent alleles A*03:01, A*11:01, C*07:01, and C*07:02, selected on the basis of their presentation by malignant cells but not normal cells. We found tumor antigens derived from genetic dependencies in lymphomas and leukemias that bind HLA-A3, -A11, -C7 molecules. While we could detect circulating T cell responses in healthy individuals to A*02:01 and A*11:01 phosphopeptides, we did not find such responses to A*03:01 or C*07:01 phosphopeptides, except when utilizing allogeneic donor T cells, indicating that these phosphopeptides may not be immunogenic in an autologous setting but can still be targeted by other means.

**How this study might affect research, practice or policy** – An expanded patient population expressing alleles other than A*02:01 can be addressed through the development of immunotherapies specific for phosphopeptides profiled in the present work, provided the nuances we describe between alleles are taken into consideration.

## BACKGROUND

T cells recognizing tumor-selective antigens presented in the context of MHC are capable of inducing durable regressions of cancers refractory to standard treatment when adoptively transferred into patients. Adequate selection of antigen targets is critical to induce remission and prevent relapse. Neoantigens produced by nonsynonymous somatic mutations represent a tumor-exclusive class of antigens but are typically private to each patient and more likely to be present when tumor mutational burden is sufficiently high. Differentially expressed tumor-associated antigens, such as WT1, Survivin, PRAME, and NYESO1 have been safely targeted to treat a variety of pediatric and hematologic tumors[1–3], which generally harbor too few mutations to produce adequate neoantigens. A significant pitfall in antigen selection is insufficient peptide presentation in that tumor antigen epitopes that are found to be immunogenic are not necessarily endogenously presented by tumors. Advances in mass spectrometry (MS) solve this problem by enabling direct sequencing of the immunopeptidome by identifying peptides eluted from HLA immunoprecipitates (HLA-IP)[4]. HLA-IP followed by MS of cell lines and primary samples has enabled identification of post-translationally modified peptides, such as phosphopeptides[5] and glycopeptides[6] as an emerging class of tumor antigens.

Several phosphopeptides have been shown to be presented by certain HLA class I and II alleles that exhibit enhanced immunogenicity, a feature hypothesized to be due to unique structural features[7–9]. Interestingly, Cobbold et al. have shown that while healthy donors harbor T cell responses specific to these phosphopeptides, leukemic patients lack such responses[10]. Moreover, these responses are restored post-allogeneic stem cell transplant.

Colorectal cancer patients have also been found to harbor TILs that recognize phosphopeptides, as well as peripheral T cells that recognize phosphopeptides at higher frequencies than healthy donors[11], closely mimicking studies of neoantigens produced by somatic mutations[12, 13].

We sought to expand the known landscape of phosphopeptides presented by prevalent HLA molecules that could be useful as immunotherapy targets. We applied HLA-IP to 10 hematologic cell line samples. We then combined our data with public immunopeptidomics datasets to assemble an expanded dataset of phosphopeptides not presented by normal tissues. From this dataset, we selected phosphopeptides of interest presented by HLA-A*03:01, - A*11:01, and -C*07:01 and studied their binding stability in vitro, in silico structural properties, and ability to stimulate T cells, to identify candidates for future immunotherapeutic treatment strategies.

## METHODS

### Cells

T2 cells, EBV-transformed B cells (EBV-BLCL), and monoclonal EBV-associated lymphoma cells (EBV-LPD) emerging post-marrow allograft were maintained in RPMI+10% FBS, supplemented 2 mM L-Glutamine, and penicillin-streptomycin. T2 cells expressing HLA alleles were generated as follows: HLA-A0301, HLA-C0701, and HLA-C0702 encoding cDNA (IDT) were inserted into pSBbi-GP[14] (Addgene plasmid # 60511) using the NEBbuilder HiFi Assembly Master Mix. Successful ligation was confirmed by Sanger sequencing of DH5a (NEB) colonies transformed with the ligation reaction. Stable transfection was performed by transfecting T2 cells with 4.5ug of HLA cDNA in the pSBbi-GP backbone and 0.5ug of pCMV(CAT)T7-SB100[15] (Addgene plasmid # 34879), followed by puromycin selection.

### HLA ligand identification

HLA ligands were isolated and identified using immunoprecipitation and liquid chromatography-tandem mass spectrometry (LC-MS/MS) as described previously[16]. Approximately 1-2x10^8^ cells were washed in PBS, snap-frozen, and stored at -80C. Pellets were thawed on ice and lysed in 1% 3-([3-cholamidopropyl] dimethylammonio)-1-propanesulfonate (CHAPS) in PBS supplemented with Roche cOmplete protease inhibitor cocktail and Roche PhosSTOP for 1 hour at 4C. Lysates were cleared by centrifugation. Supernatants were circulated over W6/32- conjugated sepharose columns for MHC class I isolation and either L243- or IVA12-conjugated sepharose columns for MHC class II isolation using a peristaltic pump overnight at 4C. Peptide-HLA complexes were eluted from dried columns in 1% trifluoroacetic acid (TFA). Peptide-HLA complexes were adsorbed onto Sep-Pak tC18 columns (Waters) pre-equilibrated with 80% acetonitrile (ACN). Peptides were eluted in 40% ACN 0.1% TFA or 30% ACN 0.1% TFA. Solid-phase extraction of peptide eluates was performed using in-house C18 minicolumns (Empore) washed with 80% ACN/0.1% TFA and pre-equilibrated with 1% TFA. Peptide eluates were run through the C18 minicolumn, which was subsequently washed twice with 1% TFA, and desalted peptides were eluted with 80% ACN/0.1% TFA. Samples were analyzed using a Lumos Fusion operated in data-dependent acquisition (DDA) mode. Peptides were separated using a 12cm built-in-emitter column using a 70min gradient (2-30%B, B: 80% ACN/0.1% Formic acid). 3uL of 8uL were injected. 3+ & 4+ (and undetermined charge states) peptides were allowed in the mass range: m/z 250-700. 2+ peptides were selected in the mass range m/z 350- 1000 while 1+ peptides were selected in the range: m/z 750-1800.

### T2 stabilization

T2 cells expressing the indicated HLA molecule were harvested from culture and incubated for 18h at room temperature. Cells were subsequently washed with PBS and resuspended in serum- free RPMI with 3ug/mL Beta-2-microglobulin (MP Biomedicals) and the indicated concentration of peptide for 3 hours at room temperature followed by 3 hours at 37C in 5% CO2. Cells were washed, stained for 30 minutes at 4°C with FVD Violet (1:1000), HLA-A2 PE (BD, 1:100), and HLA-A3 PE-Vio770 (Miltenyi, 1:100) or HLA-C AF647 (BioLegend, 1:100). Stained cells were washed twice and acquired on a BD LSRII.

### Molecular docking

Peptide docking to HLA molecules was performed as described previously[17]. Solved crystal structures of peptide-HLA complexes were retrieved from the PDB and used as template structures. PDB entry 5VGE was used to dock 9-mers to HLA-C0702, and 3RL1 and 3RL2 were used for docking 9- and 10-mers, respectively, to HLA-A0301. To generate a template for HLA- C0701, UCSF Chimera[18] was used to incorporate the K66N and S99Y mutations that distinguish C0701 from C0702. Peptides of interest were threaded onto the template by mutating the peptide using the Dunbrack and SwissSideChain rotamer libraries[19, 20] implemented in UCSF Chimera. Structures were prepacked and docked using the FlexPepDock protocol in refinement mode in Rosetta3[21, 22]. For each distinct peptide-HLA complex, each complex was scored in Rosetta energy units (REU) using the Rosetta3 full-atom score function ref2015, and the top 10 lowest REU models were selected among 200 high-resolution models. UCSF Chimera was used to visualize models and analyze hydrogen bonds.

### Immunogenicity assessment

Phosphopeptides were assessed for their immunogenicity by ELISpot analysis or multimer staining of sensitized HLA-typed donor PBMC. PBMC were sensitized to phosphopeptides selected for each donor’s HLA typing using a method described previously[4]. For dendritic cell (DC) priming, the method of Wölfl & Greenberg[23] was used. In some cases, autologous peptide-pulsed CD14+ cells or T2 cells expressing the relevant HLA allele, were used instead of DC. On day 10-13 after initial sensitization or priming, cells were restimulated with peptide-pulsed, lethally irradiated autologous PBMC, autologous DC, or T2 cells and thereafter maintained in media containing IL7/15 at 5ng/mL and IL2 (Miltenyi) at 50 IU/mL. All cultures were maintained in Xvivo-15 media(Lonza)+5% human AB serum(Gemini). Cells were typically analyzed on day 10-13 of each stimulation cycle via multimer staining or ELISPOT against autologous peptide-pulsed targets as described previously[24–26].

Dextramers were assembled according to our protocol[27], based on a previous method[28]. For dextramer enrichment from PBMC, 1-3x10^8^ PBMC were incubated in 50nM dasatinib and FcR block for 30 minutes at 37C, then incubated with 10ug/mL of the indicated dextramer pool for 1 hour at 4C, and enriched with anti-Cy5, -PE, or -Cy7 beads (Miltenyi) sequentially over 2 LS columns (Miltenyi). Enriched cells were either stained for surface markers and analyzed by flow cytometry or expanded in 96-well plates with irradiated allogeneic feeders and anti-CD3/28 reagent (STEMCELL) in the presence of IL2 300IU/mL, IL7 5ng/mL, and IL15 5ng/mL for 10-14 days until subsequent analysis or further enrichment and re-expansion. When dextramer-enriched samples were acquired by flow cytometry, the entire sample volume was acquired.

### Data analysis and statistics

Flow cytometry data was analyzed in FCS Express (De Novo Software). All other data were analyzed using custom Python and R scripts. All statistical analyses presented used a paired t-test. For genetic dependency analysis, data were downloaded from DepMap portal[29](www.depmap.org/). For each group of cell lines, DEMETER2 scores were averaged for each gene using a custom Python script. Gene ontology (GO) analysis was performed using STRING(www.string-db.org). Peptide predictions were performed using NetMHCpan4.0.

Peptide motif analysis was performed using GibbsCluster-2.0. MS data from previous studies[4,30–32] were obtained on ProteomeXchange (accession nos: PXD004746, PXD005704, PXD012083, PXD013831).

## RESULTS

### Identification of immunogenic phosphopeptides presented in the malignant state

We attempted to elucidate the phosphopeptidome for common HLA alleles other than A*02:01 to define tumor antigens presented by malignant cells, but not by normal tissue. Because of the documented presentation of phosphopeptides on EBV-BLCL, we used HLA class I and II-based immunoprecipitation and LC-MS/MS to isolate HLA ligands from 6 EBV-BLCL lines as well as 2 AML lines treated with either decitabine to enhance antigen presentation or DMSO, 1 EBV- LPD line, and 1 healthy B cell sample. By restricting the resultant peptide identifications to ≥8- mer peptides filtered by a stringent FDR of 1% and DeltaMod ≥20 in the Byonic environment[33], we recovered 40,557 unique non-phosphopeptides and 255 unique phosphopeptides across all samples. HLA class I peptidomes contained 214 unique phosphopeptides derived from 194 human proteins, whereas class II peptidomes contained 53 unique phosphopeptides derived from 37 proteins. EBV-transformed samples, such as EBV-BLCL and EBV-LPD, were the top-ranking samples when comparing samples by counts of unique unmodified peptides and phosphopeptides in our dataset (Fig. 1A). HLA class I and II phosphopeptides were most frequently 9-mers and 16-mers (Fig. 1B), respectively, in agreement with previous studies[8, 32]. To elucidate the differences in phosphopeptidome of B cells in the malignant vs. healthy state in an autologous setting, we examined class I peptides and phosphopeptides eluted from equal numbers of EBV-BLCL and healthy B cells from the same donor (donor AP). When constrained to peptides with a NetMHCpan4-predicted percentile rank ≤2% for each donor-expressed allele, high affinity peptides were more numerous for the HLA- A*11:01 allele than other allotypes present in Donor AP, with the majority of A*11:01-predicted peptides under the common 500nM cut off (dashed lines in Fig. 1C). We found more than twice the number of predicted high affinity peptides presented by donor AP’s EBV-BLCL compared to their autologous healthy B cells (5,789 vs. 2,142 peptides; Fig. 1C). Moreover, phosphopeptides, demarcated by grey circles in fig. 1C, were presented in higher numbers by each allele in EBV- BLCL compared to autologous healthy B cells (enumerated in fig. 1D). On the basis of their amino acid sequences excluding phosphorylation, most of these phosphopeptides were assigned to HLA-A*11:01 by NetMHCpan4.0. By analyzing the overlaps in phosphopeptidomes of all 6 HLA-A*11:01+ samples in our dataset (Fig. 1E), we found previously undescribed phosphopeptides pGTF3C2, pPPP1R12A, pPIM1, pMYBBP1A, and pSRRM1 were presented by multiple EBV-BLCL and EBV-LPD, but not healthy B cells. Since some phosphopeptides have been shown to be immunogenic in A2+ donors[10,17,34], we sought to determine if A11+ donors also harbor phosphopeptide-specific T cells. We selected phosphopeptides and normal peptides presented by donor AP’s EBV-BLCL, and found that only pGTF3C2 elicited peptide- specific IFNg production by sensitized T cells from this donor (Fig. 1F). Similar to previous reports, the unmodified peptide, GTF3C2wt, did not induce specific T cells. These results were corroborated by tetramer staining of donor AP’s PBMC after a single stimulation with pGTF3C2 (Fig. 1G). To explain the difference in immunogenicity, we used a molecular docking approach that we previously applied to explain the specificity of a TCR-mimicking antibody recognizing the A2/pIRS2 phosphopeptide[17]. We docked pGTF3C2 and GTF3C2wt to HLA-A*11:01 using FlexPepDock[21], and examined the top 10 lowest energy models for each complex (Fig. 1H). Both models exhibited similar mean energy scores and peptide backbone conformations, consistent with the observation that GTF3C2wt is predicted to be a strong binder, exhibiting a 0.19% percentile rank for binding to HLA-A*11:01, and that GTF3C2wt was co-presented with pGTF3C2 in our MS data. Still, pGTF3C2 exhibited more solvent facing character due to phosphorylated P4Ser (Fig.1H, left). Since T cell receptors typically strongly recognize P4-P6 of a peptide, these docking results are concordant with the increased immunogenicity we observed for pGTF3C2. These results demonstrated that certain phosphopeptides were presented by HLA- A*11:01 exclusively in the malignant state and can mobilize phosphopeptide-specific T cell responses in normal donors even when the wildtype peptide is co-presented by HLA-A*11:01.

**Figure 1:**
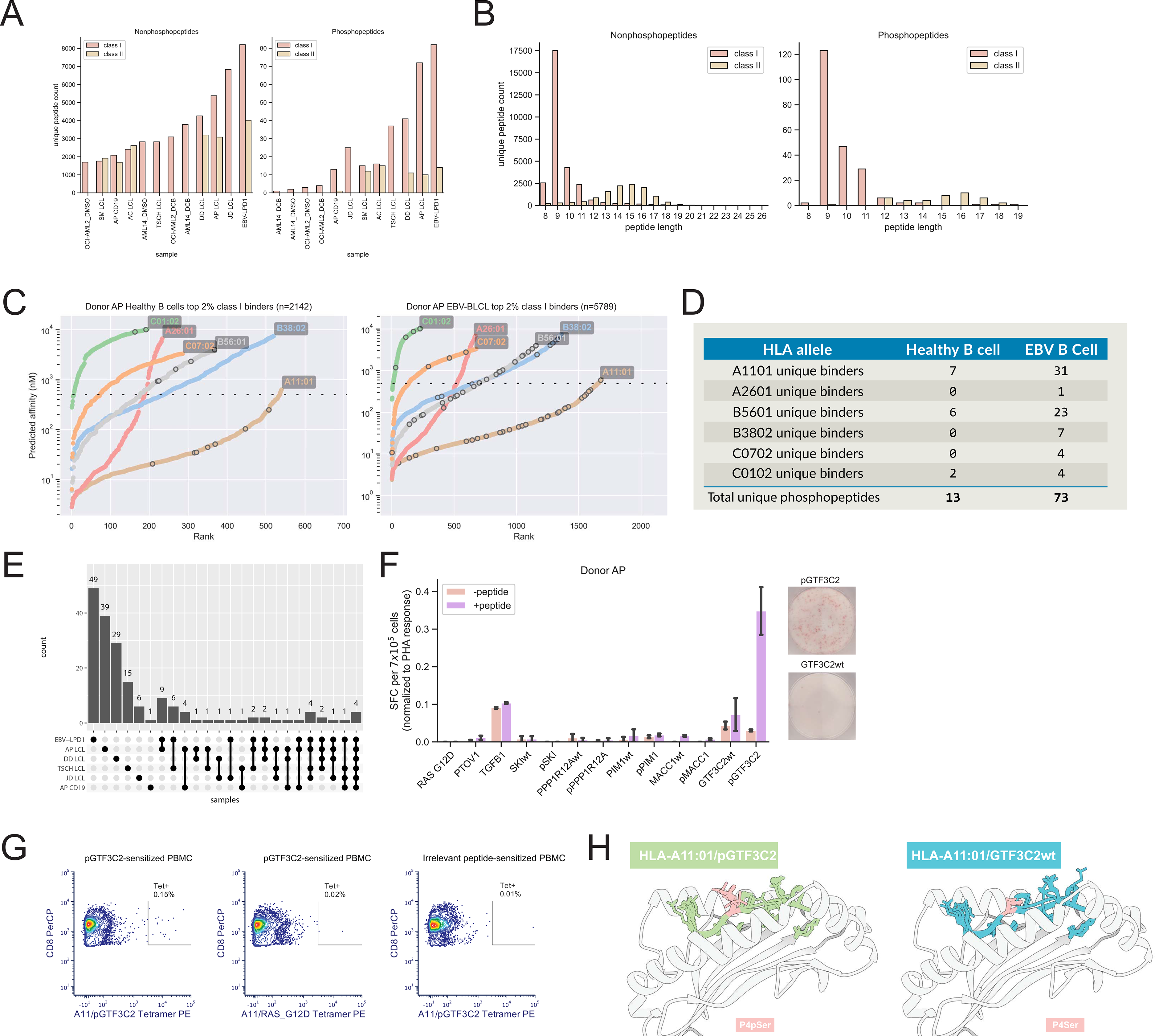
The A*11:01 immunopeptidome contains recurringly presented, immunogenic phosphopeptides. (A) Unique peptide counts for class I and II nonphosphopeptides (left) and phosphopeptides (right) across all samples on which we performed HLA-IP. (B) Length distribution of class I and II nonphosphopeptides (left) and phosphopeptides (right). (C) NetMHCpan4.0 predicted affinity versus peptide affinity rank for each HLA allotype present in healthy B cells (left) and EBV-BLCL (right) from the same donor (Donor AP). Grey circles indicate the position of phosphopeptides. Dashed line indicates 500nM affinity. (D) Total counts of unique phosphopeptides between Donor AP’s autologous healthy B cells and EBV-BLCL visualized in C. (E) UpSet analysis of overlap in phosphopeptides between all HLA-A1101+ samples on which we performed HLA-IP. (F) ELISpot of Donor AP PBMC sensitized to the indicated peptides, expressed as % of PHA-stimulated control for each peptide-sensitized culture. Sensitized cultures were restimulated on ELISpot plate with either peptide pulsed (+peptide) or unpulsed (-peptide) autologous PBMC. Representative images for pGTF3C2- and GTF3C2wt-sensitized PBMC are shown to the right. (G) Tetramer staining of Donor AP PBMC showing increased frequency of pGTF3C2 tetramer-specific CD8 T cells after pGTF3C2 sensitization relative to both irrelevant tetramer (A11/RAS_G12D) stained cells and irrelevant peptide-sensitized autologous PBMC. (H) Docking results of the top 10 lowest energy models of pGTF3C2 (left) or GTF3C2wt (right) in complex with HLA-A*11:01, with p4Ser highlighted to show effect of phosphoserine on increasing solvent-facing character.

### An expanded phosphopeptide dataset yields shared HLA-A3 supertype phosphopeptide tumor antigens

We further expanded our dataset to include phosphopeptides curated from previous studies[4,30– 32], yielding a dataset containing 2,466 distinct phosphopeptides spanning 20 tissue types, predominantly acute myeloid leukemia (n=19), mantle cell lymphoma (n=19), meningioma (n=18), and EBV-BLCL (n=13) (Fig. 2A). To extend our studies of the A*11:01 allele, we focused on phosphopeptides presented by HLA-A*03 supertype alleles, such as A*03:01 and A*11:77. We found 772 phosphopeptides presented by A3+ or A11+ samples but not by samples expressing other alleles and compared their presentation across all samples to find recurringly presented phosphopeptides (Fig. 2B). Hierarchical clustering of the samples presenting these phosphopeptides did not necessarily correlate with their tissue of origin or cancer type; however, such an analysis is limited by the uneven distribution of tissues and HLA alleles in the dataset.

**Figure 2:**
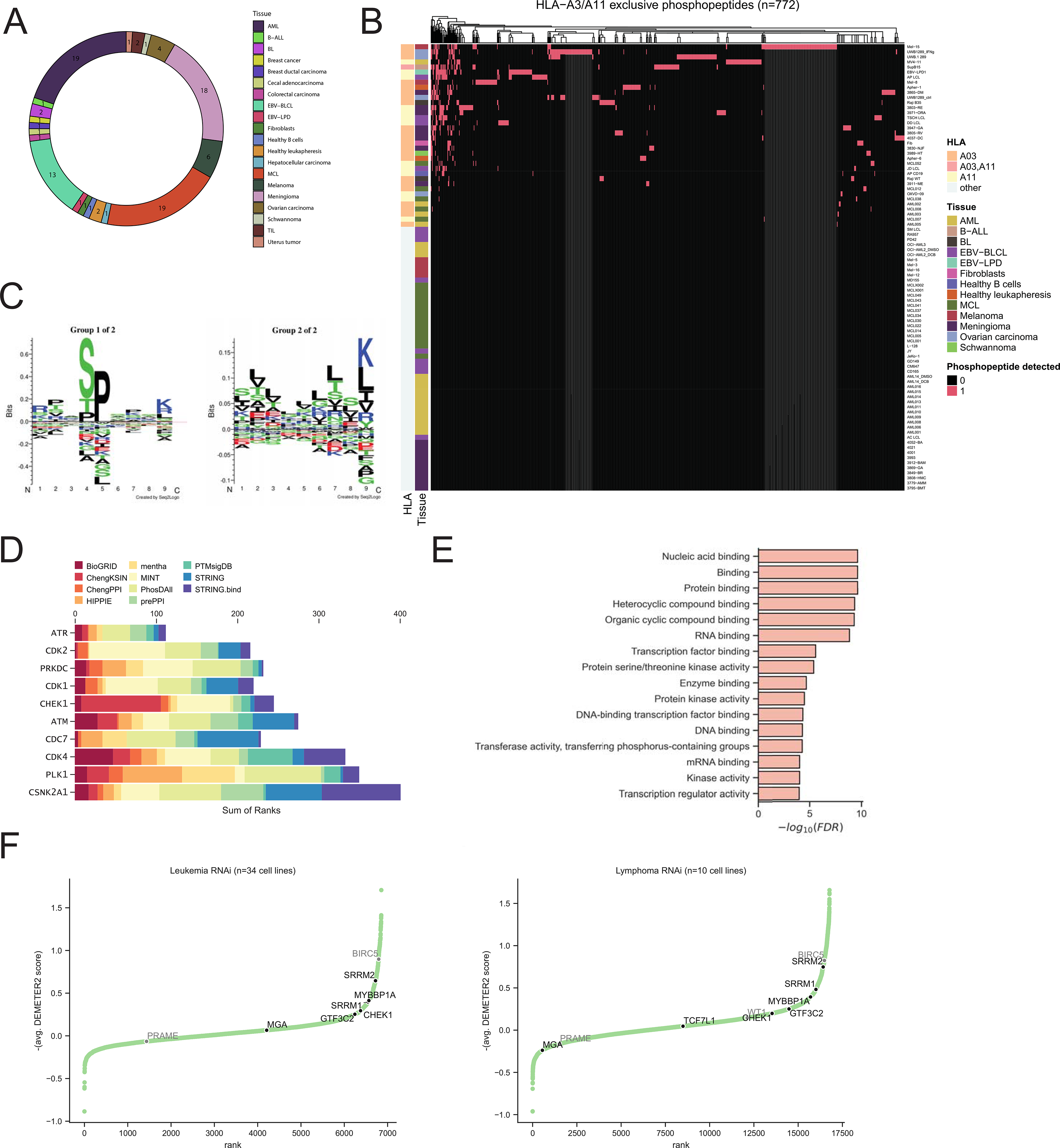
Expanded dataset of phosphopeptides. (A) Circle plot of tissue types represented in expanded dataset showing counts of unique samples of each tissue type. (B) Heatmap visualization of HLA-A3/A11 phosphopeptide presentation. Vertical axis represents distinct tissue samples. Horizontal axis represents a distinct phosphopeptide. Samples are annotated on the left by HLA allotype and tissue type. (C) Motif analysis of phosphopeptides in B. (D) Kinases inferred to be upstream of parental genes of the phosphopeptides visualized in B. Kinases are ranked by their MeanRank, with MeanRank decreasing from bottom to top, and plotted against the sum of their ranks in each of the kinase libraries, which are indicated by color. (E) GO term analysis of parental genes of phosphopeptides in (B). (F) Rank plots of genetic dependency score (-1*average (DEMETER2 score)) for lymphoma and leukemia cell lines from pooled RNAi screens[29] with parental genes of phosphopeptides denoted. Genetic dependency increases from bottom to top. Known tumor-associated antigens Survivin (BIRC5) and WT1 are denoted in grey for reference.

Unsupervised clustering of the 719 ≥9-mer A3/A11 phosphopeptide sequences revealed two motifs: 68% belong to a motif dominated by P4 Ser and P5 Pro, a preference consistently reported for phosphopeptides[4,10,32]; 32% belong to a motif characterized by repeated leucines, reflecting the representation of A*03:01 samples in the dataset since submotifs of A*03:01 contain an overrepresentation of leucine[35] (Fig. 2C). Recurringly presented A3/A11 phosphopeptides are summarized in Table 1. The majority of these phosphopeptides were detected on both A3+ and A11+ samples. To infer the upstream kinase pathways involved in phosphorylation, we performed kinase enrichment analysis[36] of the parental genes for all phosphopeptides presented by at least two malignant A3/A11 samples, but not by healthy samples. This analysis ranked the essential kinases ATR, CDK2, and PRKDC as most probable upstream kinases, all of which are involved in cell cycle and DNA repair. Kinases are plotted by decreasing MeanRank versus sum of ranks in each kinase library in Fig. 2D. Gene ontology analysis revealed that our phosphopeptides preferentially derive from nucleic-acid binding proteins (Fig. 2E). Moreover, HLA-A3/A11 phosphopeptides that were recurringly presented derived from genetic dependencies in leukemia and lymphoma cells, such as SRRM1, SRRM2, GTF3C2, and MYBBP1A (Fig. 2F). That the HLA-A3/A11 phosphopeptidome samples peptides from proteins essential to lymphoma and leukemia cell proliferation supports the categorization of selected phosphopeptides as shared tumor antigens.

**Table 1:** Summary of selected phosphopeptides from the expanded dataset presented by HLA-A3 and/or HLA-A11. * denotes phosphopeptides selected for further study

| Sequence | n<br>samples<br>presented<br>on (n<br>healthy<br>samples) | Gene | HLA | Malignant tissue presentation |
| --- | --- | --- | --- | --- |
| RVAsPTSGVK | 15 (3) | IRS2 | A03,<br>A11 | B-ALL, Melanoma, Ovarian carcinoma,<br>Meningioma, Schwannoma, MCL |
| SVSsPVKSK* | 15 (1) | MGA | A03,<br>A11 | EBV-LPD, EBV-BLCL, B-ALL, Melanoma |
| RTNsPGFQK | 12 (0) | RBM26 | A03,<br>A11 | EBV-LPD, EBV-BLCL, B-ALL, Ovarian<br>carcinoma, Meningioma, BL |
| HVYtPSTTK | 11 (3) | ANKRA2 | A03,<br>A11 | B-ALL, Melanoma, Meningioma, Schwannoma,<br>AML |
| RTAsPPPPPK* | 11 (0) | SRRM1 | A03,<br>A11 | EBV-LPD, EBV-BLCL, Melanoma, AML, MCL,<br>BL |
| KLRsPFLQK | 10 (2) | DBNL | A03,<br>A11 | B-ALL, Meningioma, AML, BL |
| KVQGsPLKK | 10 (1) | AKAP12 | A03,<br>A11 | B-ALL, Ovarian carcinoma, Meningioma,<br>Schwannoma |
| KVSsPTKPK* | 10 (1) | GTF3C2 | A03,<br>A11 | EBV-LPD, EBV-BLCL, B-ALL, Melanoma,<br>Ovarian carcinoma |
| RAKsPISLK | 10 (1) | CARD11 | A03,<br>A11 | EBV-BLCL, Meningioma, Schwannoma, MCL |
| RLSsPISKR | 10 (1) | BARD1 | A03,<br>A11 | Melanoma, Ovarian carcinoma, Meningioma,<br>AML, BL |
| ATAsPPRQK | 9 (1) | SRRM2 | A11 | EBV-LPD, EBV-BLCL, B-ALL, Meningioma |
| ATQsPISKK* | 9 (0) | MYBBP1A | A03,<br>A11 | EBV-BLCL, EBV-LPD, B-ALL, Ovarian<br>carcinoma, Meningioma |
| GSGsPAPPR | 9 (1) | GPATCH8 | A11 | EBV-LPD, EBV-BLCL, B-ALL, Meningioma |
| SVKsPVTVK | 9 (1) | TCF7L1 | A03,<br>A11 | Meningioma, Melanoma, Schwannoma |
| RTAsPNRAGK* | 3 (0) | HIF1A | A03,<br>A11 | EBV-LPD, B-ALL, Melanoma |
| RTAsPPALPK* | 4 (0) | PRDM2 | A03,<br>A11 | EBV-BLCL, EBV-LPD, Melanoma, BL |
| HSLsPGPSK* | 4 (0) | PIM1 | A11 | EBV-BLCL, Meningioma |
| ATPTSPIKK* | 8 (0) | PPP1R12A | A11 | EBV-BLCL, EBV-LPD, B-ALL, Meningioma,<br>MCL |

### Structural features of phosphopeptide complexes

Phopshopeptide-MHC complexes have been postulated to be immunogenic by virtue of their phosphate moiety conferring an increase in both MHC stabilization and solvent-facing character. However, this characteristic is a not a universal feature as structural studies of phosphopeptide- HLA-A2 complexes have shown this feature varies on a peptide-by-peptide basis. To determine if phosphorylation conferred augmented peptide binding to HLA-A3 molecules, we determined the binding of selected phosphopeptides and their unmodified (“wildtype”) counterparts to MHC via *in vitro* and *in silico* assays. Using TAP-deficient T2 cells, modified to express HLA- A*03:01, we found that in 5 of 5 phosphopeptide-wildtype pairs, both peptides could stabilize HLA-A*03:01 with no promiscuous binding to HLA-A*02:01, but the phosphorylated peptide did not confer increased stabilization of HLA-A*03:01 compared to its wildtype counterpart (Fig. 3A-C). In the case of one HLA-A3 phosphopeptide, pMGAP, phosphorylation appears to dampen the capacity of the peptide to stabilize HLA-A3, requiring a higher concentration of peptide to achieve the same degree of stabilization as unmodified MGAP, similar to what has been reported for phosphopeptides binding B*07:02 and B*40:01[32, 37]. By docking each phosphopeptide and its wildtype counterpart to HLA-A3, we found that in 2/5 phosphopeptide- wildtype pairs, MYBBP1A and MGAP, the unmodified peptide binds to HLA-A3 more stably than its respective phosphopeptide as demonstrated by lower Rosetta energy scores of the top 10 most stable complexes (Fig. 3D). Only in the case of SRRM1 did its phosphopeptide yield a significantly more stable complex than its unmodified peptide. For the two remaining phosphopeptide-unmodified pairs, HIF1A and PRDM2, there was no large difference between the stability of binding to HLA-A3. For MGAP, however, both in vitro stabilization and molecular docking suggest that phosphorylation reduces the stability of the peptide-HLA-A3 complex. To determine the differences conferred by phosphorylation of MGAP, we examined the top 10 models for pMGAP and MGAPwt and found they exhibit nearly congruent backbone conformations (Fig. 3E). The mean number of hydrogen bonds formed at the binding interface was not significantly different (11.7 for MGAPwt vs. 12.1 for pMGAP; two-tailed p-value=0.39), leading us to further investigate the interfacial interactions. Because the Rosetta score is a weighted linear combination of individual energy terms, we decomposed the Rosetta energy function into its individual terms and calculated the mean ΔΔ*G* of each term between the MGAPwt and pMGAP models to clarify the energetic favorability underlying MGAPwt. The strongest energetically favorable change was observed in electrostatic potential (Δfa_elec, -54.5 kcal/mol), which was offset by an unfavorable but smaller magnitude change in solvation energy (Δfa_sol) of 29.5 kcal/mol (Fig. 3F). Since the only difference between the pMGAP and MGAPwt structures is the presence or absence, respectively, of a phosphate moiety on P4Ser, we examined which residues on HLA interact with P4Ser to stabilize MGAPwt (Fig. 3G), finding that in MGAPwt, P4Ser interacts with Asn66 with a large mean potential of -0.574 kcal/mol.

**Figure 3:**
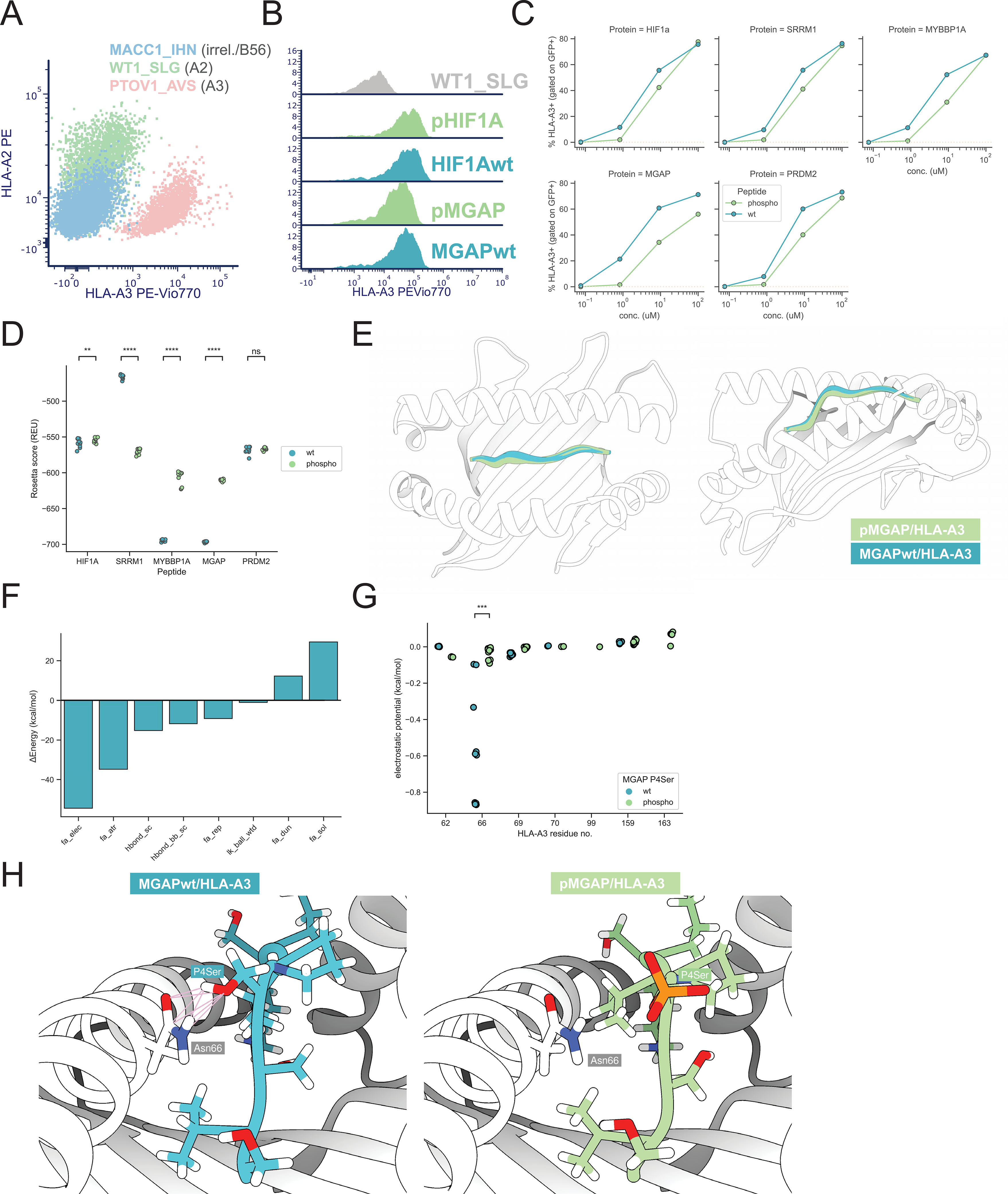
HLA-A0301 phosphopeptide binding properties. (A) Validation of HLA-A0301 stabilization using reference HLA-A3-binding peptide (PTOV1), A2-binding peptide (WT1_SLG), and irrelevant peptide (MACC1). (B) Histograms of HLA-A3 staining on T2- A0301 pulsed with 100ug/mL of the indication peptides. (C) HLA-A3 stabilization in response to dose titration of phosphopeptide-wildtype pairs. (D) Rosetta score in REU units of top 10 models produced by FlexPepDock for each phosphopeptide-wildtype in complex with HLA-A3. (E) Visualization of peptide backbone conformations of top 10 models for pMGAP and MGAPwt in complex with HLA-A3. (F) Decomposition of mean ΔΔ*G* by Rosetta energy score terms between the top 10 MGAPwt and pMGAP models visualized in (E). (G) Electrostatic potential of all P4Ser interactions with HLA-A3 for pMGAP and MGAPwt. (H) Visualization of P4Ser interacting with Asn66 in MGAPwt/HLA-A3 (left) and pMGAP/HLA-A3 (right) complexes. Pink lines indicate VDW overlaps sufficient to produce contacts between P4Ser and Asn66. All statistics produced by paired t-test. p-value annotation legend: ns: 5.00e-02 < p <= 1.00e+00; *: 1.00e-02 < p <= 5.00e-02; **: 1.00e-03 < p <= 1.00e-02; ***: 1.00e-04 < p <= 1.00e-03; ****: p <= 1.00e-04.

Visualizing P4Ser in relation to Asn66 reveals sufficient van der Waals (VDW) radii[38] overlap to constitute 7 contacts between these two residues in the MGAPwt/HLA-A3 structure (Fig. 3H, left); however, in the pMGAP/HLA-A3 structure, P4Ser does not achieve sufficient proximity to Asn66 to make such contacts (Fig. 3H, right). Nevertheless, since 3/5 phosphopeptide-wildtype pairs that we examined were co-presented in our MS data, our results demonstrate that HLA-A3 can capably present both phosphopeptides and their wildtype counterparts.

Given the limited polymorphism of the HLA-C locus relative to the -A and -B loci, we sought to characterize phosphopeptides presented by prevalent HLA-C alleles as potential tumor antigens. To this end, we selected phosphopeptides found on multiple HLA-C*0701 and -C*0702 expressing EBV-BLCL, tumors, and monoallelic cell lines[39], excluding phosphopeptides whose sequences were detected in healthy cadaver tissue[40]. We selected 7 phosphopeptides, pRBM14, pRAF1, pMYO9B, pZNF518A, pWNK, pSTMN, and pNCOR, and pulsed them onto T2 cells, modified to express HLA-C*0701 or -C*0702 to measure their capacity to stabilize each HLA. Excepting pWNK, all of the selected phosphopeptides were found on both C0701+ and C0702+ cell line immunopeptidomes. In spite of this, none of them stabilized C*0702, but all of them stabilized C*0701 on T2 cells (Fig. 4A). To investigate the molecular determinants of binding to HLA-C7, we docked each 9-mer phosphopeptide onto either C*0701 or C*0702. For each phosphopeptide complex, the top 10 models were consistently more stable for C*0701 than C*0702 phosphopeptide complexes, as measured by a lower Rosetta energy score (fig. 4B). Despite the significant energetic difference between the complexes, the peptide backbone conformations were similar between C*0701 and C*0702 complexes, with C*0701-bound phosphopeptides being slightly more shifted out of the HLA groove (Fig. 4C); however, the buried interfacial solvent-accessible surface area (SASA) was only significantly different between pNCOR-bound C*0701 and C*0702 complexes (Fig. 4D). Examining representative structures for pNCOR shows that the discrepancies of conformations between C7 molecules is mediated by different sidechain orientations (Fig. 4E). Since C*0701 and C*0702 differ by only two residues at positions 66 and 99, we hypothesized that direct contacts with these residues may explain the difference in conformations and interfacial surface area. pNCOR makes 12 and 13 hydrogen bonds to C*0702 and C*0701, respectively, but few of these bonds are mutually observed in both complexes. When pNCOR is bound to C*0701, the sidechain of P2Arg makes two hydrogen bonds with Tyr99 and one with Asp9, both belonging to the β sheet of the HLA B-pocket. When bound to C*0702, which bears a serine at position 99 instead of tyrosine, P2Arg did not form a hydrogen bond with the distant Ser99 molecule (Fig. 4F), thereby losing two hydrogen bonds in the B-pocket β sheet. This change resulted in C*0702-bound pNCOR forming additional hydrogen bonds with the HLA α1 helix between the sidechain of P1Arg and Glu63, and between the phosphate moiety of P4pSer and Arg69. No direct contacts were observed between the other distinguishing residue at position 66. These changes in the C*0702 phosphopeptide complex resulted in a peptide conformation that is more buried in the HLA groove and less energetically favorable. These results suggest altered B-pocket interactions underlie the stability of phosphopeptides when bound to C*0701. Only 2/7 of these phosphopeptides, pRAF1 and pWNK, were found to be co-presented with their wildtype counterpart in our MS data. Docking their phosphopeptide-wildtype pairs to C0701 revealed no energetic differences or peptide backbone differences (data not shown). The fact that 5/7 selected phosphopeptides differentially expressed by tumors bound C0701 without co-presentation of their wildtype peptide further suggests an abundance of phosphopeptide presentation by C0701, concordant with previous observations[32].

**Figure 4:**
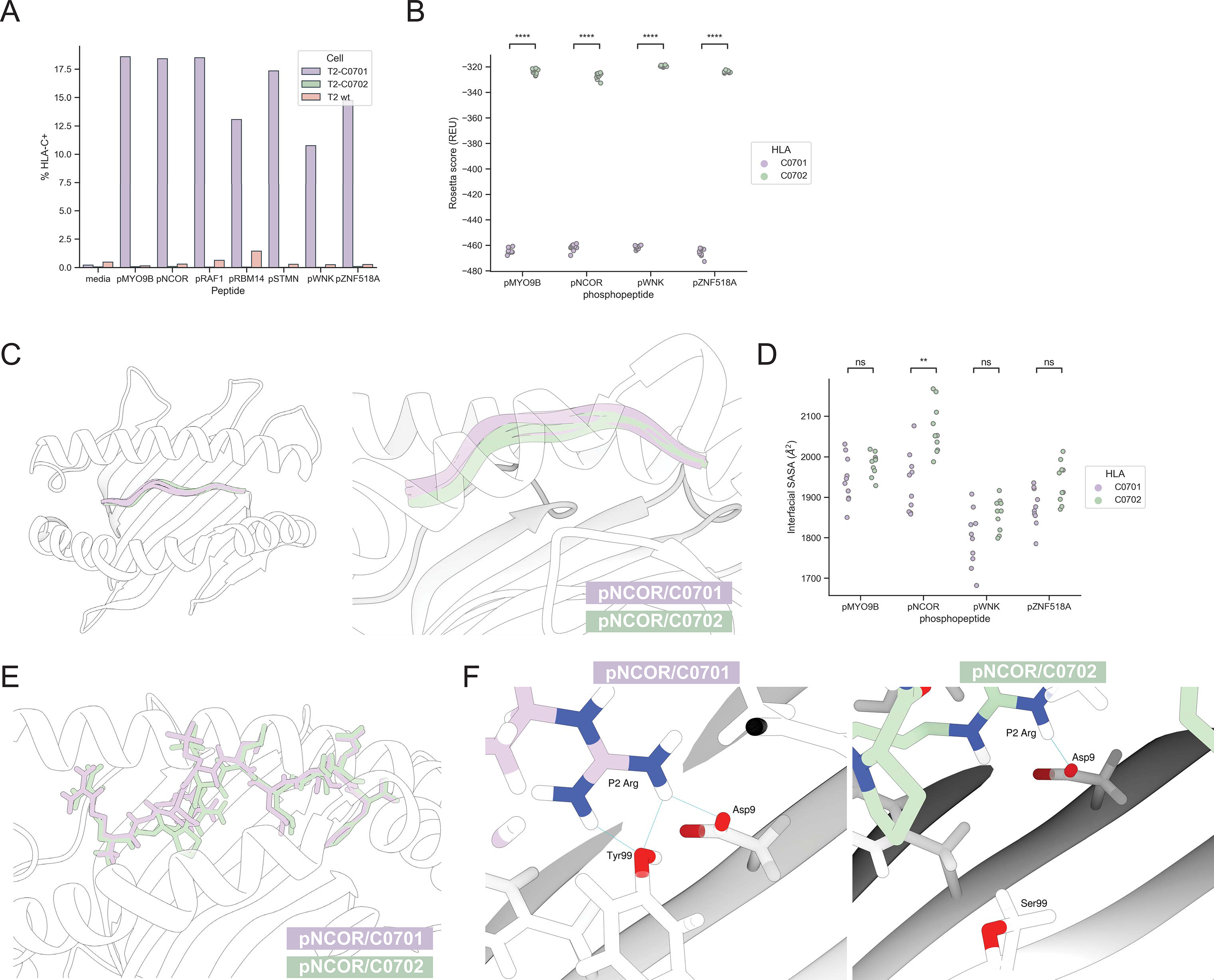
HLA-C7 phosphopeptide binding properties. (A) HLA-C stabilization of indicated phosphopeptides at 100ug/mL for T2-C0701, -C0702 or wt. (B) Rosetta score of top 10 models produced by FlexPepDock for each indicated phosphopeptide in complex with HLA-C0701 and - C0702. (C) Visualization of peptide backbone conformation for the top 10 models for each of pNCOR/C0701 and pNCOR/C0702. (D) Buried interfacial solvent accessible surface area (SASA) at the interface between each indicated phosphopeptide and C0701 or C0702. (E) Representative visualization of sidechains for pNCOR/C0701 and pNCOR/C0702. (F) Visualization of hydrogen bonds formed between P2Arg and Tyr99 in the pNCOR/C0701 complex (top) and absence of hydrogen bonds between P2Arg and Ser99 in pNCOR/C0702 (bottom). All statistics produced by paired t-test. p-value annotation legend: ns: 5.00e-02 < p <= 1.00e+00; *: 1.00e-02 < p <= 5.00e-02; **: 1.00e-03 < p <= 1.00e-02; ***: 1.00e-04 < p <= 1.00e-03; ****: p <= 1.00e-04.

### Immunogenicity assessment

The immunogenicity of HLA-A*0201-presented phosphopeptides in healthy donors has been previously shown to derive from memory, rather than naïve, T cells, unlike other responses to self-antigens [10, 34]. Given the prevalence of phosphopeptide-specific T cells (PP-CTL), we sought to detect PP-CTL via phosphopeptide dextramers assembled using an improved method[28]. We assembled HLA-A*0201 dextramers in complex with 6 selected phosphopeptides as well as an irrelevant A2-binding WT1 peptide, WT1_SLG, and used fluorophore barcoding[41] to discriminate phosphopeptide specificity from WT1 specificity (Fig. 5A). We stimulated A*02:01^+^ donor PBMC with A*02:01^+^ T2 cells pulsed with the selected 6 phosphopeptides and found that after only 10 days, 4% phosphopeptide dextramer^+^ CD8^+^ T cells could be detected that did not cross-react with WT1_SLG dextramer. However, PBMC from the same donor contemporaneously stimulated with T2 cells pulsed with the A2-binding WT1_SLG peptide did not yield T cells specific for either WT1_SLG or phosphopeptides (Fig. 5B), providing evidence that the immunogenicity of phosphopeptides is more pronounced than that of WT1_SLG despite our observation that these phosphopeptides and WT1_SLG comparably stabilize HLA-A2. In two additional A*02:01+ healthy donor buffy coats, we were able to use sequential magnetic enrichment to achieve a highly pure (>95% dextramer^+^) PP-CTL population (Fig. 5C). To determine if PP-CTL could exert effector function, we measured the specific IFNg response of PP-CTL by ELISpot. Across 4 A*02:01+ donors, responses could be detected to pIRS2 and pCDC25B (Fig. 5D,E), the latter of which were consistently phosphorylation- specific.

**Figure 5:**
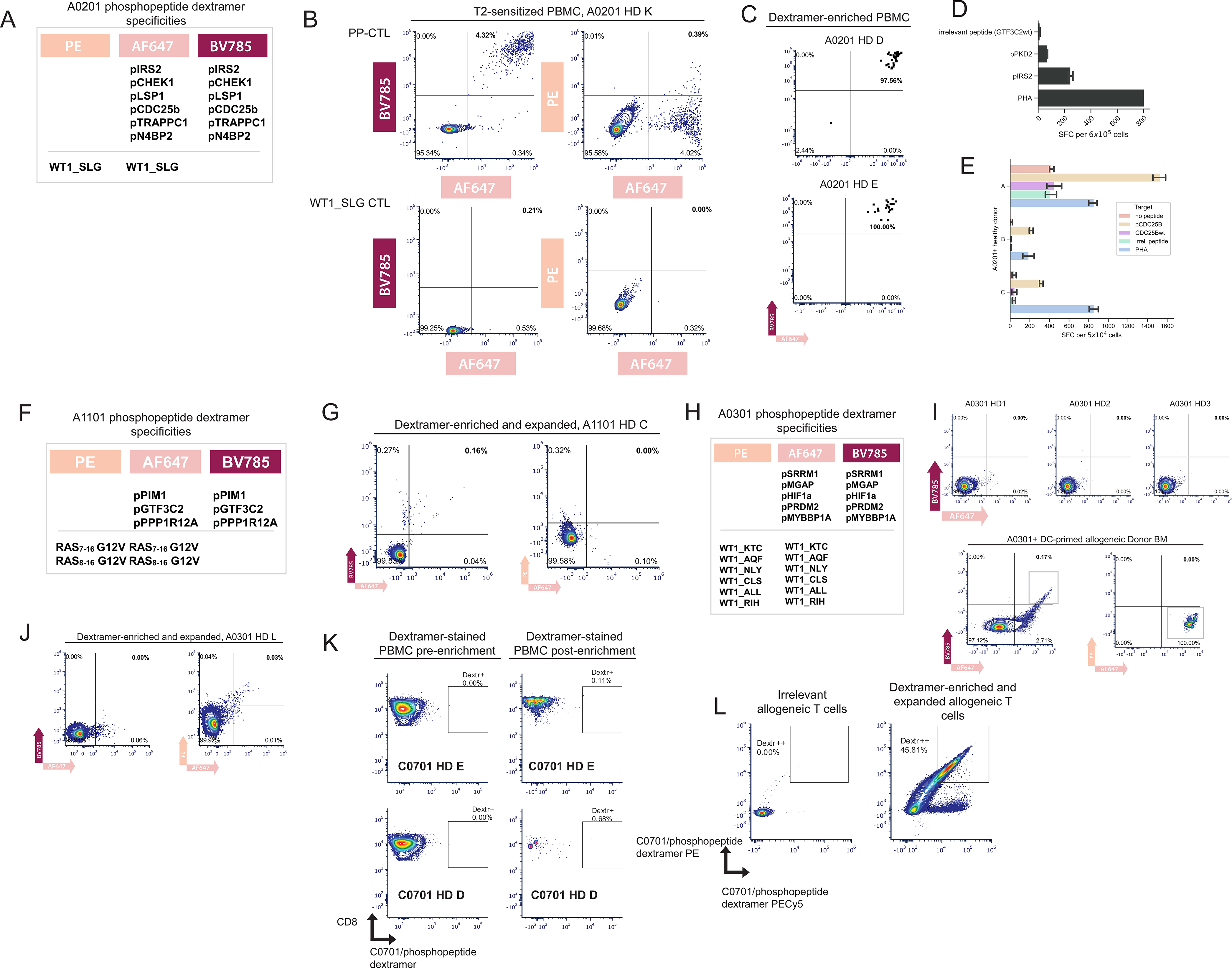
Combinatorial fluorophore barcoded dextramer analysis of PP-CTL. (A) HLA-A0201 dextramer panel. HLA-A2/phosphopeptide dextramer is encoded by AF647/BV785 and A2/WT1_SLG is encoded by PE/AF647 combination. (B) A0201+ donor T cells sensitized to either phosphopeptides (top) or WT1_SLG (bottom). PP-CTL (top) can be detected by A2/phosphopeptide dextramers in the AF647/BV785 channel and do not crossreact to A2/WT1_SLG dextramer in the PE/AF647 channel. (C) Analysis of A0201+ donor buffy coats enriched for PP-CTL by sequential dextramer enrichment over 2 magnetic columns and then analyzed by flow cytometry. (D) ELISPOT of A0201 healthy donor T cells sensitized to A2 phosphopeptides pPKD2 and pIRS2. PHA response is shown as positive control and irrelevant GTF3C2wt peptide response serves as negative control. (E) ELISPOT of 3 A0201 donors (denoted A, B, C on y-axis) whose T cells were repeatedly sensitized to pCDC25B and then rechallenged with autologous CD14+ cells pulsed with either of the indicated peptides listed under “Target”. PHA serves as a positive control. (F) A1101 dextramer panel encoding phosphopeptides on BV785/AF647 and RAS G12V peptides on PE/AF647. (G) A1101 Donor PBMC enriched with dextramers and expanded show binding to A1101/phosphopeptide dextramers (AF647/BV785) but not irrelevant RAS G12V dextramers (PE/AF647). (H) HLA- A0301 dextramer panel encoding phosphopeptides on AF647/BV785 multiple A3-binding WT1 epitopes on PE/AF647. (I) Three A0301+ donors primed to the phosphopeptides in H. did not result in expansion of A3/phosphopeptide dextramer-binding cells (top), but A0301-negative donor BMMC stimulated with A0301+ DC produce A3/phosphopeptide dextramer-binding T cells that do not crossreact with A3/WT1 dextramer. (J) Dextramer-enrichment and expansion increases frequency of A3/WT1-specific T cells, but not PP-CTL. (K) C0701 Healthy donors do not have directly observable PP-CTL after dextramer enrichment with C0701/phosphopeptide dextramers on a single fluorochrome. (L) Dextramer-enriched and expanded T cells from C0701- negative donors bind C0701/phosphopeptide dextramer on dual fluorochromes (PE/PECy5). All flow plots gated on live, CD8+ CD19-CD14-CD123-CD40- single cells.

Since we observed responses to certain A*11:01 phosphopeptides, we assembled an A*11:01 dextramer panel encoding either phosphopeptide or irrelevant RAS G12V specificity (Fig. 5F). When these dextramers were used to enrich and expand PBMC from an A*11:01+ healthy donor for PP-CTL, a small population of PP-CTL could be observed that did not crossreact to irrelevant RAS G12V peptide dextramers (Fig. 5G). Unlike our observations of A*02:01 PP-CTL, we did not observe A*11:01 PP-CTL directly after sequential magnetic enrichment of PBMC from 2 additional A*11:01+ healthy donor buffy coats (data not shown). Since our enrichment scheme co-enriches for RAS G12V specificity, our data suggest that A*11:01 phosphopeptide responses are at least more prevalent than those of RAS G12V, but still not as prominent as responses to A*02:01 phosphopeptides.

To detect HLA-A*03:01 PP-CTL, we assembled A*03:01 dextramers using shared phosphopeptides identified from our analysis denoted in Table 1 and used A3-binding epitopes of WT1 as irrelevant controls (Fig. 5H). We could not detect PP-CTL directly ex vivo after dextramer enrichment of A*03:01 PBMC (data not shown), nor could we observe PP-CTL after priming with phosphopeptide-loaded autologous DCs in 3 A*03:01+ healthy donors (Fig. 5G) or repeated sensitization with phosphopeptide-pulsed T2-A0301 (data not shown). However, after priming A*03:01-negative donor BMMC-derived T cells with allogeneic phosphopeptide-loaded A0301+ DC, we could observe PP-CTL that bound A3 phosphopeptide dextramer (Fig. 5H, top), but not irrelevant A3 WT1 dextramer (Fig. 5H, bottom). Interestingly, when we used a co-enrichment scheme specific for the shared WT1/phosphopeptide fluorochrome AF647 and expanded the resulting T cells, a minor population of WT1-specific, rather than phosphopeptide-specific, T cells was observed (Fig. 5J). Overall, A0301 phosphopeptides appear to be less immunogenic than A*03:01 WT1 peptides, and A*11:01 and A*02:01 phosphopeptides, requiring, in our hands, stimulation of A*03:01-negative allogeneic T cells with loaded A*03:01 targets to elicit a phosphopeptide-specific response.

To determine if there were any immediate responses to phosphopeptides binding C*07:01 that we previously described, we assembled a dextramer pool containing pRBM14, pRAF1, pMYO9B, pZNF518A, pWNK, pSTMN, and pNCOR in complex with C*07:01. In two C*07:01 healthy donor buffy coats we could not detect responses prior to or after sequential dextramer enrichment (Fig. 5K); however, using iterative dextramer enrichment and expansion cycles, we could generate a population of strongly dextramer-binding T cells from a C*07:01-negative donor (Fig. 5L). These data suggest that these C*07:01 phosphopeptides, although in high abundance in the immunopeptidome[32], do not generate prevalent T cell responses in the autologous setting similar to A*03:01 phosphopeptides.

## DISCUSSION

Phosphopeptides represent an emerging class of HLA ligands that have potential to be targeted as shared tumor antigens produced not by mutations, but by cancer associated post-translational modifications. Since T cell responses to phosphopeptides have been studied comprehensively in the context of HLA-A*02:01 and -B*07:02[10,34,42], we sought to expand the phosphopeptidome by systematically reanalyzing large immunopeptidomics datasets containing phosphopeptides, yielding a new set of phosphopeptides presented by the HLA-A3 supertype as well as C*07:01. We further used healthy tissue databases[40] to determine the extent to which these phosphopeptides are found on healthy tissue. Our observation that there were phosphopeptides whose HLA binding adheres to HLA-A3 supertype classifications while being presented exclusively by malignant cells supports the potential of phosphopeptides as tumor antigens that could be targeted across multiple alleles. Supertype binding has been observed previously for phosphopeptides, such as in the case of the same epitope of pIRS2 binding to A*02:01 and A*68:02, and its length variant binding A*03:01 and A*11:01[43]. Functionally although aberrant phosphorylation is considered a hallmark of cancer, the phosphopeptides we identified are mostly derived from common essential pathways, such as nucleic acid binding and repair, rather than oncogenic kinase pathways. Interestingly, differential presentation of phosphopeptides has been ascribed more to inhibition of critical phosphatases than kinase overactivation[43].

We used TAP-deficient cells to verify that MS-identified phosphopeptides could stabilize their cognate HLA alleles. All phosphopeptides selected for A*03:01 and C*07:01 bound their cognate allele; however, A*03:01 phosphopeptides did not exhibit a binding advantage compared to their wildtype counterparts. Previously, the half-lives and IC50 values of phosphopeptide-HLA complexes were found to be improved over wildtype counterparts for only 1/5 HLA-A*01:01 peptides, 0/5 HLA-B*07:02 peptides, and 0/7 HLA-B*40:02 peptides[32, 37]. This is in contrast to studies of HLA-A2, where phosphorylation was found to increase the binding affinity of a given peptide by 1.1–158.6-fold in 10/11 cases[9]. However, it must also be recognized that the discrepancies in binding properties of phosphopeptide-wildtype pairs between HLA alleles may be affected by technical limitations. Specifically, MS data acquired in data-dependent acquisition (DDA) mode will contain increased representation of charged peptides, which fragment more efficiently. Since HLA-A2 alleles prefer hydrophobic anchor residues, MS-identified HLA-A2 phosphopeptides may be more likely to contain suboptimal anchor residues that preclude HLA binding in the absence of phosphorylation. This technical limitation may be the underlying cause for identification of phosphopeptide-wildtype pairs that are co-presented by HLA-A3, which prefers charged anchor residues. Interestingly, in phosphopeptide-wildtype pairs, phosphopeptides more frequently displayed enhanced binding to HLA-C*07:02 and -C*06:02 compared to -A and -B alleles[32]. In our data, we found that most of our selected C*07:01 phosphopeptides were not co-presented with their wildtype counterparts, but the two phosphopeptide-wildtype pairs that were presented did not display any differences in energetics based on molecular docking, suggesting that while co-presentation can occur, the phosphopeptide is the more abundantly presented of the pair. These data highlight the importance of considering the residue preference of the HLA allele of interest when selecting phosphopeptides with augmented binding properties.

Our structural studies based on molecular docking determined that peptide sidechain interactions with the HLA B-pocket and α1 helix were critical determinants of phosphopeptide binding to HLA-A3 and -C7. Previously, HLA-A0201 phosphopeptide binding was shown to be dependent on phosphate-mediated contacts made with Arg65 of the α1 helix[9]. A more recent study found that in HLA-B0702 phosphopeptide complexes, the phosphate moiety was within H-bond distance to Arg62 of the α1 helix[44]. In the case of HLA-A0301, we found that a wildtype P4Ser could mediate more favorable contacts with Asn66 of the α1 helix than phosphoserine.

However, the absence of these interactions observed in HLA-A3/pMGAP attenuated binding. In a more dramatic case, we compared phosphopeptide binding to HLA-C*07:01 and -C*07:02, alleles which only differ by two B pocket residues at 66 and 99[45]. We found that Tyr99 of C0701 forms hydrogen bonds with P2 that were absent due to the presence of Ser99 of C0702.

These results demonstrated that B-pocket and α1 helix interactions can shape the character of peptide binding to HLA in the presence of subtle differences like phosphate modification or residue substitutions. Our results obtained by molecular docking are qualified, however, by the absence of solvent interactions, which are accounted for in x-ray crystallography and explicit-solvent molecular dynamics.

A feature of phosphopeptides that motivates development of phosphopeptide-targeted agents is that the phosphorylated epitope sequences present a different recognition surface than their wildtype counterparts. Thus, several studies have shown that T cells, TCRs, or TCR-like antibodies specific for MHC-presented phosphopeptides specifically recognize phosphate moieties without crossreactivity to wildtype peptides[10,17,44,46]. In contrast, class II-restricted pWED-specific T cells did not differentiate between phosphopeptide and wildtype counterpart[47]. Therefore, TCR-like agents capable of phosphorylation discrimination can be developed, but such discrimination is not guaranteed when evaluating native T cells, necessitating the use of adequate methodologic procedures to generate phosphorylation-specific candidates.

Our study demonstrates that phosphopeptides are potentially shared tumor antigens, but nuances between alleles present important considerations for the development of cancer immunotherapies. Namely, A*03:01 and B*07:02 are similar in their co-presentation capacity of phosphopeptide-wildtype pairs, whereas A*02:01 generally exhibits a binding preference for phosphopeptides. However, A*02:01 and B*07:02 phosphopeptides are more frequently immunogenic in an autologous setting than A*03:01 phosphopeptides. Therefore, A3 phosphopeptide targeting may require the use of allogeneic or synthetic TCRs to redirect otherwise absent T cells, or the use of TCR-mimicking antibodies to directly engage immune effectors.

## DECLARATIONS

### Ethics approval and consent to participate

Written informed consent was received from participants prior to inclusion in the study. Use of human blood samples was approved by MSKCC IRB protocol #06-107.

### Consent for publication

All donors provided consent for publication of de-identified data in this study where applicable.

### Availability of data and material

Original datasets (mass spectrometry search results, flow cytometry data, docking results) will be made available by the authors upon reasonable request to the corresponding author (Zaki Molvi, Memorial Sloan Kettering Cancer Center, New York, NY, USA)

### Authors’ Contributions

All authors made substantial contributions to the study. ZM conceived and designed the study, developed methodology, acquired data, analyzed and interpreted data, and wrote the original draft of the manuscript. MGK and TD developed methodology, acquired data, and analyzed and interpreted data. JU acquired data. ZM, MGK, TD, JU, DAS, and RJO reviewed and revised the manuscript. DAS and RJO supervised the study.

## Acknowledgments

We thank the Dr. Henrik Molina of the Proteomics Resource Center at Rockefeller University for the performance of all LC/MS-MS experiments. Molecular graphics and analyses performed with UCSF Chimera, developed by the Resource for Biocomputing, Visualization, and Informatics at the UCSF, with support from NIH P41-GM103311. We acknowledge the use of the MSK Flow Cytometry Core Facility, funded in part through the NIH/NCI Cancer Center Support Grant P30 CA008748. We thank Dr. Joshua Elias (Chan Zuckerberg Biohub, Stanford, CA, USA) for providing HLA typing of samples from PXD004746 and PXD005704.

## LIST OF ABBREVIATIONS

MHC: major histocompatibility complex
HLA: human leukocyte antigen
MS: mass spectrometry
HLA-IP: HLA immunoprecipitation
LC-MS/MS: liquid chromatography-tandem mass spectrometry
CHAPS: 3-([3-cholamidopropyl] dimethylammonio)-1-propanesulfonate ACN: acetonitrile
TFA: trifluoroacetic acid
EBV-BLCL: Epstein-Barr virus transformed B lymphoblastoid cell line
EBV-LPD: Epstein-Barr virus-associated B lymphoproliferative disease
AML: acute myeloid leukemia
B-A LL: B cell acute lymphoid leukemia
MCL: mantle cell lymphoma
TIL: tumor infiltrating lymphocytes
PP-CTL: phosphopeptide-specific T cells
PBMC: peripheral blood mononuclear cells
VDW: van der Waals
DDA: data-dependent acquisition

